# Sfp1 regulates transcriptional networks driving cell growth and division through multiple promoter binding modes

**DOI:** 10.1101/420794

**Authors:** Benjamin Albert, Susanna Tomassetti, Yvonne Gloor, Daniel Dilg, Stefano Mattarocci, Slawomir Kubik, David Shore

## Abstract

Understanding how transcriptional programs help to coordinate cell growth and division is an important unresolved problem. Here we report that the nutrient-and stress-regulated transcription factor Sfp1 is rate-limiting for expression of several large classes of genes involved in yeast cell growth, including ribosomal protein, ribosome biogenesis, and snoRNA genes. Remarkably, the spectrum of Sfp1 transcription effects is concordant with a combination of chromatin immunoprecipitation and chromatin endogenous cleavage binding analyses, which together provide evidence for two distinct modes of Sfp1 promoter binding, one requiring a co-factor and the other a specific DNA-recognition motif. In addition to growth-related genes, Sfp1 binds to and regulates the promoters of cell cycle “START” regulon genes, including the key G1/S cyclins *CLN1* and *CLN2*. Our findings suggest that Sfp1 acts as a master regulator of cell growth and cell size by coordinating the expression of genes implicated in mass accumulation and cell division.

## Introduction

The expression of genes required for ribosome production is an intensive transcriptional process in growing cells (Warner, 1999) and serves as a paradigm to study coordination of large gene networks (Lempiainen & Shore, 2009). Regulation of ribosome production at the transcriptional level in eukaryotes is best understood in the budding yeast *Saccharomyces cerevisiae*, where RNA polymerase II (RNAPII)-mediated transcription of ribosomal protein (RP) genes, the suite of >200 protein-coding genes required for ribosome assembly (referred to as ribosome biogenesis [RiBi] genes), and small nucleolar RNA (snoRNA) genes is highly coordinated and regulated according to nutrient availability and stress. Despite this fact, the promoters of these three groups of genes are organized differently, begging the question of how they can be coordinately regulated (Bosio, Negri et al., 2011).

The Split-Finger Protein 1 (Sfp1) (Blumberg & Silver, 1991) is a nutrient-and stress-sensitive transcription factor (TF) that has emerged as a potential coordinator of cell growth and division. Deletion or over-expression of *SFP1* influences expression of a large number of genes related to growth, including RP and RiBi genes (Fingerman, Nagaraj et al., 2003, Jorgensen, Rupes et al., 2004, Marion, Regev et al., 2004). Consistent with a direct role in cell growth, Sfp1 is concentrated in the nucleus under optimal growth conditions, but rapidly relocates to the cytoplasm in response to nutrient deprivation or other stress conditions (Jorgensen et al., 2004, Marion et al., 2004). In addition to its role in cell growth, cellular levels of Sfp1 also influence cell size and cell-cycle progression (Cipollina, Alberghina et al., 2005, Jorgensen, Nishikawa et al., 2002, Xu & Norris, 1998). Thus, *sfp1Δ* cells are amongst the smallest viable single-gene deletion mutants, whereas *SFP1* overexpression leads to a large-cell phenotype (Jorgensen et al., 2002). Taken together, these findings suggest that Sfp1 might play a key role in coordinating cell growth and cell division. Interestingly, the transcriptional and cell-size phenotypes of *SFP1* are notably similar to those of the c-Myc proto-oncogene (Jorgensen et al., 2004, Jorgensen & Tyers, 2004, Lempiainen & Shore, 2009).

One paradox that has limited our understanding of Sfp1’s mechanism of action is that the protein has been detected by Chromatin Immuno-Precipitation (ChIP) at only a small fraction of the promoters that it appears to regulate. For example, although ChIP detects Sfp1 at many RP gene promoters (Reja, Vinayachandran et al., 2015), it is undetectable at virtually all of the >200 RiBi gene promoters where over-expression studies suggest that it might be a direct activator (Jorgensen et al., 2002, Jorgensen et al., 2004).

Here we vastly expand our knowledge of Sfp1 binding by Chromatin Endogenous Cleavage (ChEC)-seq analysis (Schmid, Durussel et al., 2004, Zentner, Kasinathan et al., 2015). Remarkably, we find that ChEC and ChIP provide a highly complementary picture of Sfp1 binding, with distinct sets of sites identified by one technique or the other. Our combined analysis provides evidence that Sfp1 directly orchestrates TATA-binding protein (TBP) and RNAPII recruitment at a broad array of genes that drive cell growth, including most RiBi, RP and snoRNA genes. In addition, we find that Sfp1 binds to the promoters of many G1/S (“START”) regulon genes that are targeted by the TF Swi4. Interestingly, Sfp1 binding sites identified by ChEC are enriched for the motif gAAAATTTTc, whereas binding identified by ChIP is often strongly dependent on another TF: Ifh1 at RP genes or Swi4 at G1/S regulon genes. These findings provide an unprecedented example of how the combination of ChIP and ChEC can reveal a more complete picture of TF-chromatin interactions. Taken together, our results support a role for Sfp1 as a master regulator that helps to orchestrate cell growth by coordinating transcriptional programs involved in mass accumulation and cell division.

## Results

### Modulation of Sfp1 protein level triggers a genome-wide redistribution of RNAPII

Steady-state mRNA measurements in strains deleted for *SFP1* have revealed up-or down-regulation of more than 2000 genes (Cipollina et al., 2005, Cipollina, van den Brink et al., 2008, Jorgensen et al., 2004). However, *sfp1Δ* cells grow very slowly, making it difficult to distinguish between direct and indirect effects (O’Duibhir, Lijnzaad et al., 2014). Furthermore, measurements of steady-state mRNA levels can mask transcription effects that are buffered by compensatory mRNA stability changes (Sun, Schwalb et al., 2012). Therefore, to understand better the role of Sfp1 we decided to use RNAPII occupancy measured by ChIP as a read-out for transcription, first examining the effect of Sfp1 overexpression. We placed *SFP1* under the control of a strong inducible promoter (p*GAL1*) and measured RNAPII recruitment by ChIP-seq of the Ser5-phosphorylated form of RNAPII after 1h of galactose induction. Sfp1 overexpression triggered a massive change in the transcriptional program, consistent with previous findings (Jorgensen et al., 2004), with 745 genes up-regulated and 1429 genes apparently down-regulated by at least 1.5-fold (**Figure 1A;** see **Table S1** for a complete list).

We were struck by the fact that many of the genes down-regulated upon Sfp1 overexpression are glucose-repressed genes implicated in carbohydrate metabolism, whereas induced genes are strongly enriched in RP and RiBi genes, as well as translation-related genes and genes associated with “non-coding RNA metabolic processes” (see **Table S1** for GO term analysis). This global change in the transcriptional program appears similar to that observed following glucose addition to cells growing on less optimal carbon sources. To assess this resemblance more directly, we performed ChIP-seq of RNAPII 10 minutes after a glucose pulse. We found a strong overlap between genes that are repressed or activated in both conditions, including RiBi and RP genes (**Figure 1B** and **Figure S1A**). Consistent with this finding, motifs identified in the promoters of genes up-regulated by Sfp1 over-expression are highly similar to those up-regulated following a glucose pulse (**Figure S1B**). These data show that Sfp1 levels can influence expression of more than one third of RNAPII-transcribed genes, suggesting that Sfp1 could play a key role in a much larger transcriptional network than is revealed by ChIP analysis of its binding sites (Reja et al., 2015).

**Figure 1:**
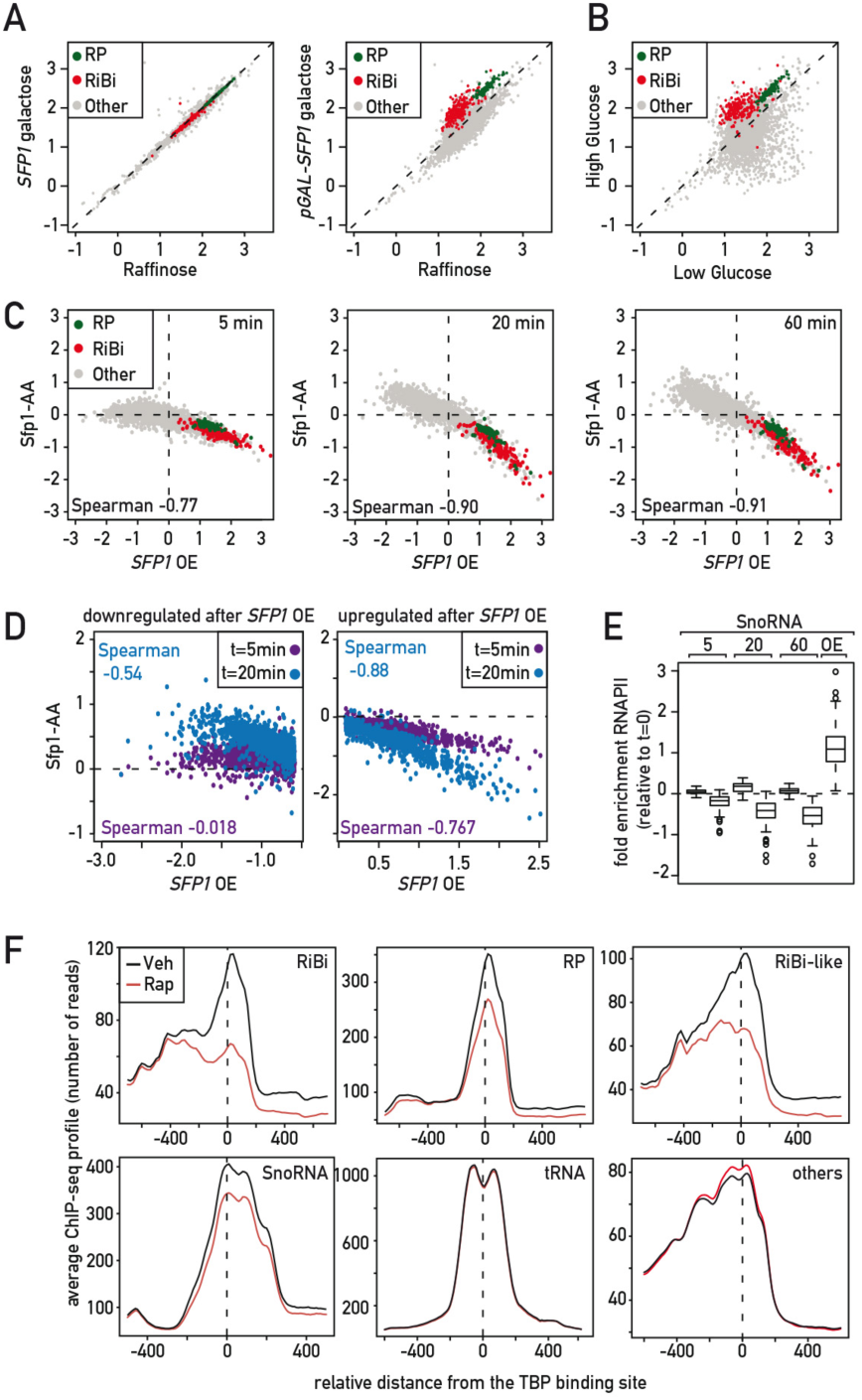
Regulation of growth-related genes by Sfp1. (A) Scatter plot comparing Rpb1 ChIP-Seq signal (log10 normalized read counts) in SFP1 (left panel) and pGAL1-SFP1 (right panel) strains grown in 2% raffinose (x-axis) or one hour following 2% galactose addition (y-axis). RP genes are indicated in green, RiBi genes in red and all other genes in grey. (B) Scatter plot, as in (A), comparing Rpb1 ChIP-Seq signal in low glucose (0.5%; x-axis) and 10 minutes after glucose addition to 2% (y-axis). (C) Scatter plots comparing Rpb1 ChIP-seq fold change (log2) relative to t=0 in a Sfp1-FRB anchor-away strain 5 (left panel), 20 (middle panel) and 60 (right panel) min following rapamycin addition (y axes) to Rpb1 ChIP-seq change relative to t=0 at 60 min following galactose addition in a pGAL-SFP1 strain (x-axes). RP and RiBi genes indicated as in (A). (D) Scatter plots derived from data shown in (C) in which genes down-regulated (left panel) and up-regulated (right panel) under conditions of SFP1 overexpression (x-axis) are compared to the effects at 5 min (purple) and 20 min (blue) following initiation of Sfp1 nuclear depletion (rapamycin addition). The Spearman correlation is indicated for each of the four separate categories. (E) Box plots showing Rpb1 ChIP-seq fold-change (log2) at snoRNA genes relative to t=0 for cells treated for 5, 20 or 60 min with rapamycin (red) or vehicle (grey) in a Sfp1 anchor-away strain (Sfp1-FRB), or 60 minutes following galactose addition to pGAL1-SFP1 cells (SFP1 OE). (F) Average TBP ChIP-seq signal centred on the TBP binding site at promoters of RiBi, RP, RiBi-like, snoRNA, tRNA, and all other genes (as indicated) 20 minutes following rapamycin (red) or vehicle (black) treatment of an Sfp1-FRB anchor-away strain.

To challenge this idea, we used the “anchor-away” system (Haruki, Nishikawa et al., 2008) to measure the immediate effect of rapid Sfp1 nuclear depletion on RNAPII association genome-wide. As expected, nuclear depletion of Sfp1 causes a growth defect (**Figure S1C**). Strikingly, nuclear depletion of Sfp1 in the anchor-away strain appears complete by ∼15 minutes (**Figure S1D**), similar to what is observed in wild-type strains following stress, inactivation of TORC1, or glucose depletion (Jorgensen et al., 2004). To ascertain which genes might be direct targets of Sfp1, we measured RNAPII binding by ChIP-seq at 5, 20, and 60 minutes following rapamycin addition to the anchor-away strain and compared these data to the changes observed following Sfp1 over-expression (1 hr growth of the *pGAL-SFP1* strain in galactose; **Figure 1C**). We observed a significant anti-correlation between depletion and over-expression effects (Spearman= 0.77, 0.90. and 0.91 after 5, 20 and 60 min, respectively, of rapamycin treatment) confirming that the majority of up-regulated and down-regulated genes identified by over-expression analysis are also sensitive to a reduction of Sfp1 nuclear levels. The weaker anti-correlation at 5 minutes, compared to 20 or 60 minutes, results largely from those genes that appear to be negatively regulated by Sfp1 (**Figure 1D**), suggesting that for at least some of these genes the inhibitory effect of Sfp1 might be a secondary effect or that mechanisms by which Sfp1 directly inhibits expression might follow slower kinetics than those by which it works as an activator. Since negative regulation (direct or indirect) by Sfp1 was unanticipated, we performed a “spike-in” control (Chen, Hu et al., 2015), using *Schizosaccharomyces pombe* chromatin (Bruzzone, Grunberg et al., 2018, Hu, Petela et al., 2015), which allowed us to confirm that the increases observed in RNAPII binding following Sfp1 depletion were not due to a normalization error in the ChIP-seq analysis.

### Sfp1 promotes PIC assembly and transcription initiation at many growth-related genes

We next analyzed in more detail the molecular roles of the genes that are both up-regulated after Sfp1 overexpression and down-regulated at 5, 20, and 60 minutes of depletion by >1.5-fold, i.e. those genes where Sfp1 appears to be a direct activator. As indicated above, this group of over 500 genes is highly over-represented by RiBi (201) and RP (112) genes (**Table S2**). Although both sets of genes are down-regulated with similar kinetics following Sfp1 depletion, the magnitude of the effect is greater for RiBi genes (**Figure S1E**). Other genes in this group display kinetics and amplitude of down-regulation most similar to that of RiBi genes (**Figure S1G**), and analysis of their promoters reveals a strong enrichment for the RRPE motif, and to a lesser extent the PAC motif, both of which are common to RiBi genes (**Figure S1F**; (Bosio et al., 2011, Hughes, Estep et al., 2000)). In addition, many of these genes share several functional annotations with RiBi genes (see **Table S2** for a complete list with GO terms), and we thus refer to this group as “RiBi-like”.

A more thorough examination of the novel Sfp1 target genes within the RiBi-like group revealed three different connections to functions previously associated with Sfp1. First, we noted a strong enrichment for genes involved in nuclear transport in the RiBi-like group, consistent with the initial identification of *SFP1* based on a phenotype of altered nuclear import when present in multiple copies ((Blumberg & Silver, 1991), see **Table S2**). Second, the RiBi-like group includes all known genes encoding proteins involved in translation termination (**Table S3**), among which are the ribosome-associated Hsp70-like proteins Ssb1/2, and the termination factors Sup45 and Sup35, all of which have also been directly implicated in prion formation in yeast (Liebman & Chernoff, 2012). Curiously, Sfp1 also exists in a prion-like form [*ISP*^+^] that suppresses the phenotype of the prion-like derivative of Sup35 [*PSI*^+^], perhaps by increasing activation of genes linked to translation termination, and thus promoting translation efficiency (Matveenko, Drozdova et al., 2016, Rogoza, Goginashvili et al., 2010, Volkov, Aksenova et al., 2002). Finally, we also identified new Sfp1 target genes with regulatory functions connected to Sfp1. One of these, *MRS6*, encodes the only yeast Rab escort protein, which in addition to its essential function in secretion, interacts directly with Sfp1 and regulates its nuclear localization (Lempiainen, Uotila et al., 2009, Singh & Tyers, 2009). Another novel target of Sfp1, *TOD6*, encodes a repressor of RiBi genes (Huber, French et al., 2011, Lippman & Broach, 2009). These regulatory links point to possible feedback mechanisms that might act to fine-tune nutrient and/or stress responses.

We then asked whether Sfp1 could be involved in transcription of snoRNA genes, a distinct set of RiBi-like genes many of whose promoters are bound by Tbf1 and Reb1, two essential general regulatory factors (Bosio et al., 2011, Preti, Ribeyre et al., 2010). Transcription of most of the 78 snoRNA genes is driven by a dedicated RNAPII promoter comprising an individual transcription unit (59 genes), whereas some are grouped in operons and a few are embedded within introns of either RP or RiBi genes (Bosio et al., 2011). Notably, snoRNA genes as a whole display significant down-regulation following Sfp1 depletion and marked up-regulation upon Sfp1 over-expression, similar to that of RiBi, RP, and RiBi-like genes (**Figure 1E**).

To investigate how Sfp1 impacts transcription, we first asked whether it influences pre-initiation complex (PIC) assembly, the first step in RNAPII recruitment, by monitoring TBP binding. Indeed, rapid nuclear depletion of Sfp1 leads to a significant drop in TBP ChIP-seq signal that tracks with the RNAPII decrease (i.e. larger at RiBi and RiBi-like genes, compared to RP and snoRNA genes; **Figure 1F**). As expected, Sfp1 depletion has no effect on TBP binding at genes where RNAPII recruitment is unaffected, or at RNAPIII-transcribed tRNA genes. Since Sfp1 has been suggested to affect RNAPII processivity, particularly at RP genes (Gomez-Herreros, de Miguel-Jimenez et al., 2012), we quantified the RNAPII distribution across ORFs following Sfp1 depletion but found no change (**Figure S1H**).

### ChIP-seq reveals dynamic carbon source-related binding of Sfp1 at G1/S network genes

To determine if Sfp1 acts directly at the promoters of the genes described above we performed a ChIP-seq experiment with a strain expressing a Sfp1-TAP fusion protein from the endogenous *SFP1* locus. Given the fact that *sfp1Δ* most strongly impairs growth in medium containing glucose as carbon source, we decided to measure Sfp1 binding in three different carbon source conditions (glucose and two “poor” carbon sources, raffinose and galactose). As reported previously (Fingerman et al., 2003, Jorgensen et al., 2002, Marion et al., 2004, Reja et al., 2015), Sfp1 promoter binding at many RP genes is observed in glucose-grown cells, but few if any binding events are detected at RiBi genes under these conditions. We also observed robust Sfp1 binding at RP gene promoters in cells grown in either galactose or raffinose (**Table S4**). However, we identified ∼100 target genes in glucose-grown cells that scored negative in both galactose and raffinose when we applied a conservative cut-off for specific binding events (see **Figure 2A** for one example, **Figure S2A, Table S4**).A quantitative analysis of Sfp1 binding at promoters of these genes showed that binding is not absent in sub-optimal carbon sources but is instead decreased by about 1.5-to 3-fold compared to that in glucose (**Figure 2B).** Strikingly, we found that the group of genes where Sfp1 binding is glucose-enhanced is highly enriched in genes implicated in the G1/S cell-cycle transition, or “START” ((Bertoli, Skotheim et al., 2013); **Figure 2B, Table S4**), whose promoters are typically bound by the Swi4 activator. In contrast, genes where Sfp1 promoter binding was essentially equivalent in all carbon sources were highly enriched in Ifh1-bound RP genes.

**Figure 2:**
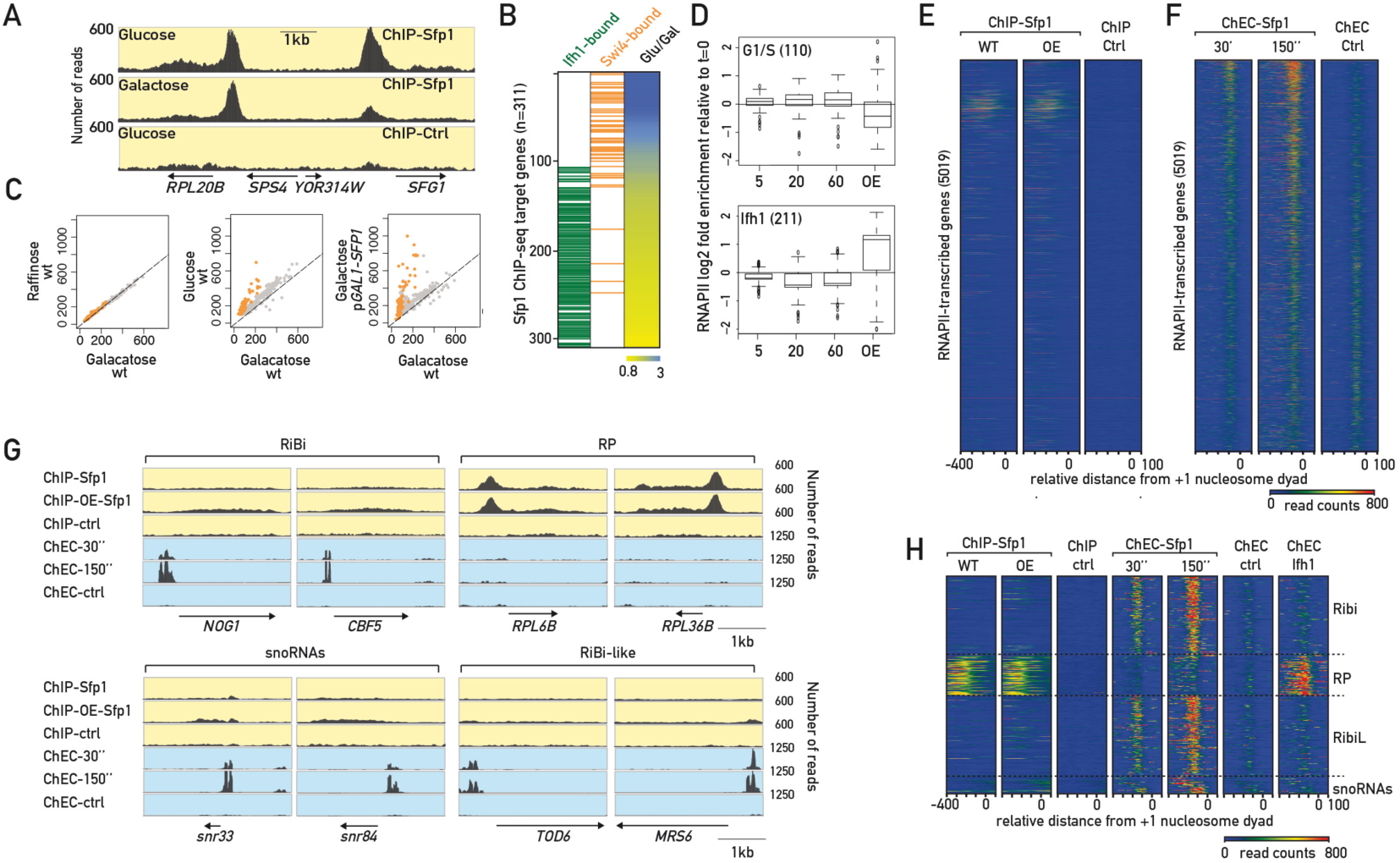
ChIP and ChEC detect distinct classes of Sfp1 promoter binding sites. (A) Genome browser tracks comparing Sfp1-TAP ChIP-seq signals in glucose or galactose medium, or in an untagged strain grown in glucose. The position and direction of transcription of individual genes is shown below. (B) Heat map showing ratio of Sfp1-TAP ChIP-seq signal from glucose-grown versus galactose-grown cells. Genes whose promoters are bound by Swi4 (Harbison, Gordon et al., 2004) or Ifh1 (Knight, Kubik et al., 2014) are indicated in orange or green, respectively. (C) Scatter plots comparing Sfp1-TAP ChIP-seq signal in a *SFP1-TAP* strain (WT) grown in galactose (x-axis) with signal in raffinose (left pane, y-axisl), glucose (middle panel, y-axis) or with a *pGAL1-SFP1-TAP* strain after 60 minutes growth in galactose (right panel, y-axis). Promoter of genes whose Sfp1 binding increases more than 1.5-fold are indicated in orange. (D) Box plots showing Rpb1 ChIP-seq fold-change (log2) at genes where Sfp1 promoter binding changes according to carbon source (“G1/S”, top panel) or where Sfp1 promoter binding is detected in all carbon sources tested (“Ifh1”, bottom panel). (E) Heat maps showing Sfp1-TAP ChIP-seq signal in a SFP1-TAP strain (WT) or a pGAL1-SFP1-TAP strain (over-expression, OE) after 60 minutes growth in galactose. ChIP-seq in an untagged strain (ctrl) is shown to the right. Rows (y-axis) consist of 5019 RNAPII-transcribed genes with a well-defined +1 nucleosome, ordered according to decreasing RNAPII fold-change after 5 minutes of rapamycin treatment in a Sfp1-FRB anchor-away strain (top to bottom). Signal densities from -400 to +100 bp relative to the +1-nucleosome dyad axis are shown (x-axis). (F) Heat maps showing Sfp1-MNase or free MNase (ctrl) ChEC-seq signal at the indicated times following calcium treatment. Genes (rows on y-axis) and signal density relative to TSS (x-axis) are as in (E). (G) Genome browser tracks comparing Sfp1-TAP or untagged ChIP-seq signals (yellow background) to Sfp1-MNase and free MNase ChEC-seq signals (blue background), the latter at the indicated time points following calcium addition. The position and direction of transcription of individual genes is shown below. (H) Heat maps showing Sfp1-TAP ChIP-seq under endogenous expression (WT) or after over-expression of Sfp1 (OE), Sfp1 ChEC-seq signal after 30 or 150 seconds of calcium treatment, and Ifh1 ChEC-seq signal after 150 seconds of calcium treatment in a window of 500 bp containing +1 nucleosome (0) at different categories of genes. Control for ChIP (untagged strain) or ChEC (free-MNase) are also shown.

To reveal if this glucose-specific increase of Sfp1 promoter binding is linked to its nuclear concentration, which is known to change according to growth conditions (Jorgensen et al., 2004), we determined whether increasing total levels of Sfp1 by growing *pGAL1-SFP1* cells in galactose could be sufficient to recapitulate the binding pattern of Sfp1 observed in glucose. Remarkably, Sfp1 overexpression specifically increased Sfp1 promoter binding at glucose-sensitive promoters but not at those binding sites common to all three carbon sources (**Figure 2C**). These data suggest that Sfp1 binding, specifically at G1/S gene network promoters, is limited by Sfp1 concentration or activity when cells are grown in the presence of a sub-optimal (non-glucose) carbon source.

To examine the function of Sfp1 at the START-specific group of genes, we quantified RNAPII association by Rbp1 ChIP-seq following both Sfp1 nuclear depletion and over-expression. In contrast to what we observed at other gene groups, Sfp1 over-expression led to a decrease in RNAPII binding at most START-specific genes, and its depletion caused a slight increase in average RNAPII binding, suggesting that Sfp1 may act as a negative regulator at many of these genes (**Figure 2D**). Interestingly, Sfp1 has been described as a negative regulator of START not only due to its ability to promote ribosome biogenesis and growth, but also through an unknown mechanism acting at the level of *CLN1/2* transcription, which drives the G1/S transition ((Aldea, Jenkins et al., 2017, Ferrezuelo, Colomina et al., 2012); see below).

### ChEC-seq reveals Sfp1 target genes that are missed by ChIP

Although the ChIP-seq experiments described above confirmed Sfp1 binding to a number of genes where functional experiments suggest it is either a positive or negative regulator, they fail to explain how Sfp1 controls expression of large groups of additional target genes, such as RiBi, RiBi-like and snoRNA genes. We thus asked whether an alternative assay to measure TF binding, chromatin endogenous cleavage (ChEC; (Schmid et al., 2004)), could reveal Sfp1 binding at the promoters of these genes. To this end, we fused the gene encoding micrococcal nuclease (MNase) to the C-terminus of the endogenous *SFP1* gene and performed a ChEC assay, results of which were analyzed by high throughput sequencing (ChEC-seq; (Zentner et al., 2015)). Strikingly, this revealed a strong signal, well above a background observed after prolonged digestion in a strain expressing free MNase, at a much larger number of promoters than was detected by Sfp1-TAP ChIP-seq (**Figure 2E, 2F, Table S5**). Significantly, target genes identified by ChEC-seq share similar functional annotations with genes that we identified above, using functional assays, as targets of Sfp1 (**Table S5**). In fact, the magnitude of the Sfp1-MNase ChEC-seq signal at promoters correlated much better than that of Sfp1-TAP ChIP-seq with the transcriptional effect observed upon Sfp1 nuclear depletion (**Figure 2E 2F, S2C**).

To examine the Sfp1 ChEC-seq results in more detail, and better compare them to those obtained by ChIP-seq, we focused on the group of over 500 genes described above, whose expression is most strongly dependent upon Sfp1. As before, we divided this group of genes into four sub-groups: the RiBi factors (as defined by Jorgensen et al. (2004)), the RP genes, the “RiBi-like” genes and the snoRNA genes. Mapping both Sfp1 ChIP-seq and Sfp1 ChEC-seq signals on these separate groups (**Figure 2H and S2D**) shows clearly that ChIP-seq reveals Sfp1 binding at RP genes, but little or no binding at RiBi, RiBi-like, or snoRNA genes. The opposite is true for ChEC-seq. Genome browser screen shots of specific examples of this effect are shown in **Figures 2G** and **S2B**.

This complementary behaviour of Sfp1 is not a universal feature of the ChEC assay as applied to TFs, since the ChEC-seq results for three general regulatory factors in yeast (Rap1, Abf1 and Reb1) are largely concordant with those obtained by ChIP (Zentner et al., 2015). We also find that Ifh1 ChEC analysis yields a profile very similar to that of ChIP (strong cleavage almost exclusively at RP genes; **Figure 2H**). Nevertheless, we have no reason to believe that the differential behaviour of Sfp1 in these two chromatin binding assays is unique to this factor.

### Co-factor dependent and sequence-driven binding modes of Sfp1

In considering possible causes for the different behaviour of Sfp1 in ChEC and ChIP assays, we first noted that most RP gene promoters, in addition to being bound by the general regulatory factor Rap1, are also bound by a highly RP gene-specific set of co-activator proteins, Fhl1 and Ifh1 (Jorgensen et al., 2004, Martin, Soulard et al., 2004, Rudra, Zhao et al., 2005, Schawalder, Kabani et al., 2004, Wade, Hall et al., 2004). In contrast, RiBi genes have not been associated with any specific activator protein(s). Since Ifh1 binding is co-incident with that of Sfp1 at RP genes (**Figure 3A**), we wondered whether Sfp1 association at these genes might be dependent on this factor. To test this idea, we measured Sfp1 binding at two RP genes following rapid nuclear depletion of Ifh1 and found that it is strongly reduced under these conditions (**Figure 3B**). This dependence upon Ifh1 for Sfp1 binding probably extends to all RP genes, since we observe a very strong correlation between the ChIP-seq strength of the two factors that is largely specific to these genes (**Figure S3A**). As noted above (**Fig. 2B**) many additional Sfp1 promoter binding sites detected by ChIP are also bound by the TF Swi4, and at these promoters we found that Sfp1 binding is highly coincident with that of Swi4 (**Figure 2C**). Anchor-away of Swi4 caused a strong decrease in Sfp1 binding at two such genes that we tested, those encoding the G1/S cyclins Cln1 and Cln2 (**Figure 3D**). We thus infer that many of the Sfp1 binding events detected by ChIP are linked to recruitment through another TF: Ifh1 at RP genes and Swi4 at G1/S regulon genes. These would appear to explain the majority of ChIP-detectable binding events, though other examples may exist where a different co-factor helps to recruit Sfp1.

**Figure 3:**
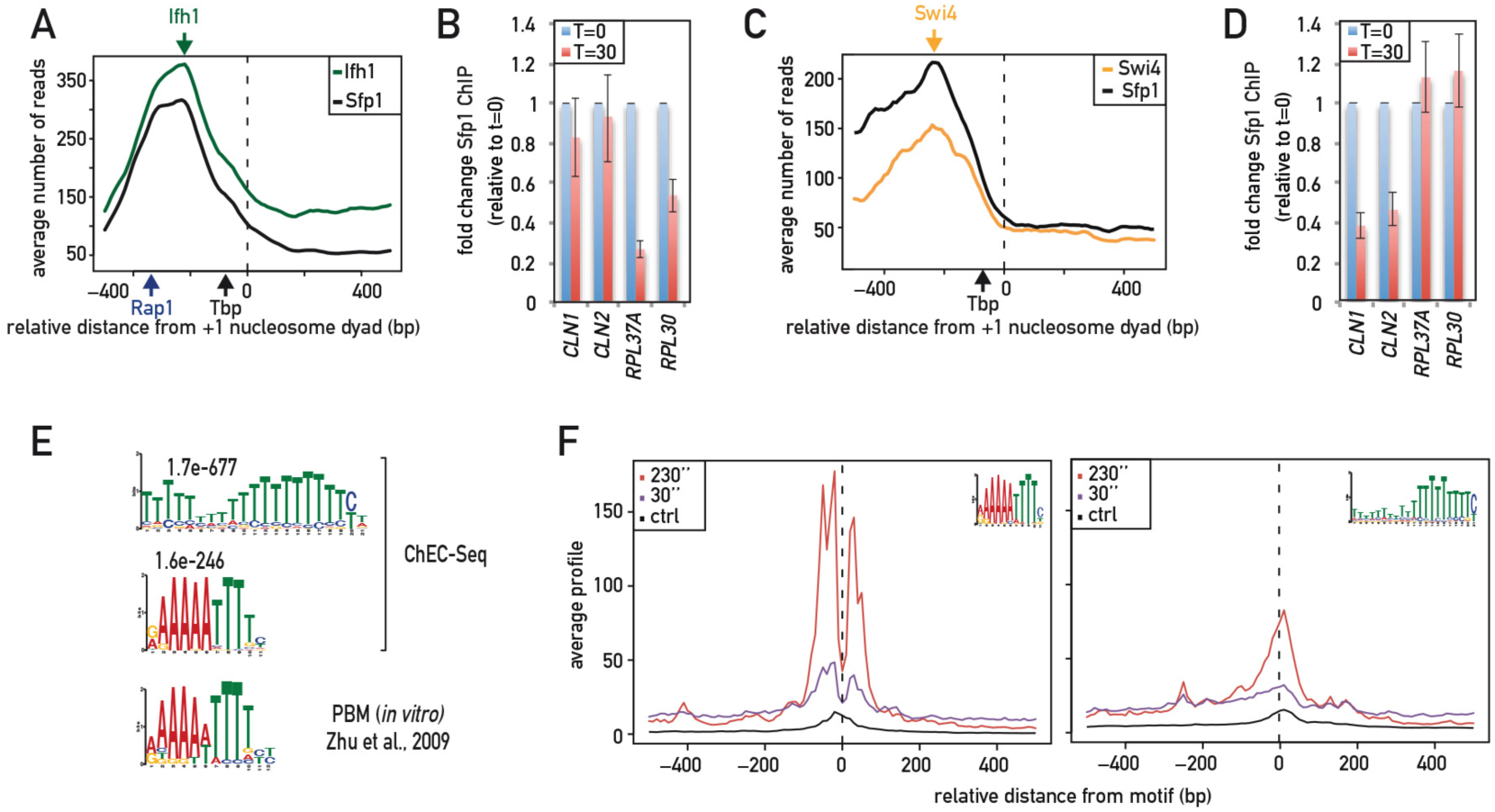
Co-factor-dependent and DNA sequence motif-dependent modes of Sfp1 binding. (A) Average ChIP-seq signal of Ifh1-Myc (green) and Sfp1-TAP (black) occupancy at RP genes, centred on the position of the +1 nucleosome. Blue and black arrows show average positions of Rap1 and TBP binding, respectively. (B) Sfp1 occupancy (qPCR-ChIP) at the *CLN1, CLN2, RPL30* and *RPL37A* promoters for the indicated times following auxin treatment in a Ifh1-AID strain; fold enrichment relative to *ACT1* was normalized to values at t = 0, which were set to 1. (C) Average ChIP-seq signal of Swi4 (green) and Sfp1-TAP (black) occupancy at Swi4 regulated genes identified in ChIP-seq of Sfp1, centred on the position of the +1 nucleosome. Black arrows show average positions of TBP binding. (D) Sfp1 occupancy (qPCR-ChIP) at the *CLN1, CLN2, RPL30* and *RPL37A* promoters at the indicated time following rapamycin treatment in a Swi4-FRB strain; fold enrichment relative to *ACT1* was normalized to values at t = 0, which were set to 1. (E) Motif enrichment identified by MEME analysis of sequences centred on Sfp1-ChEC peaks or by in vitro protein-binding microarray (PBM) analysis (Zhu, Byers et al., 2009), as indicated. (F) Average plots of Sfp1 cleavage around the indicated motifs (top right of each panel) enriched in Sfp1-ChEC (30 or 150 seconds after Ca^+2^ addition). Control averages (free-MNase cleavage 20 minutes after Ca^+2^) at these sites is also shown. Center of motif is indicated by a vertical dotted line.

To understand how Sfp1 is recruited at genes where it is detected by ChEC, we searched for a common DNA feature near the sites of Sfp1-MNase cleavage (Bailey, 2011). We found a strong enrichment for two different motifs, one a large stretch of A residues, the other a palindromic A/T-rich sequence that strongly resembles the RiBi-associated RRPE motif (**Figure 3E**). These two motifs are also enriched at promoters of genes that are affected by Sfp1 depletion or overexpression (**Figure S2B**), consistent with the high correlation of these data sets. Significantly, protein-binding microarray (PBM) data indicate that Sfp1 has DNA-binding specificity for an RRPE-like DNA sequence nearly identical to the palindromic motif identified by our ChEC experiments (Zhu et al., 2009), suggesting that Sfp1 binds directly to this motif in vivo. In contrast, the polyA motif is common to the three other yeast transcription factors characterized to date in a pioneering ChEC study (Abf1, Reb1 and Rap1; (Zentner et al., 2015)) and has been proposed to reflect a scanning mode DNA binding for these factors. We have not examined this hypothesis further but would note that the Sfp1-MNase cleavage pattern surrounding the RRPE-like motif is distinct from that seen at polyA tracts (**Figure 3F**) and suggestive of stable binding at the RRPE motif and cutting on either side of the motif, as opposed to the broad observed cleavage across the non-specific site (Zentner et al., 2015).

## Discussion

Results described here help to clarify the previously enigmatic role of Sfp1 in transcription and directly place this protein at the center of transcriptional networks controlling ribosome biogenesis and other growth-promoting processes, as well as the G1 to S transition (START) (**Figure 4**). Although previous studies indicated that Sfp1 is an activator of RiBi genes, this conclusion was based upon steady-state mRNA measurements in an extremely slow growing *sfp1Δ* strain or upon *SFP1* overexpression. The absence of a Sfp1 ChIP signal at RiBi genes thus raised serious concerns that its effect at these genes might be indirect. Our findings put these concerns to rest by demonstrating robust association of Sfp1 with RiBi gene promoters, using ChEC-seq, and by revealing that rapid and acute Sfp1 nuclear depletion,by anchor-away, results in immediate and strong down-regulation of these genes. We note at the same time, though, that many genes, most of which are weakly expressed in normal growth conditions, appear to be negatively regulated by Sfp1, since their expression increases upon Sfp1 nuclear depletion and decreases upon Sfp1 over-expression (**Fig. 1C, D**). Given that most of these Sfp1-repressed genes show no evidence of Sfp1 promoter binding (*CLN1* and *CLN2* being notable counter-examples, see **Fig. 3D** and below), how can one explain this regulation? One possibility is that the massive down-regulation of highly-transcribed genes upon Sfp1 nuclear depletion releases significant amounts of RNAPII, and/or important general co-activators, that through mass action increase the expression of many weakly transcribed genes where polymerase and co-activators might be limiting. This explanation is consistent with the delayed effect of up-regulation upon Sfp1 withdrawal but remains to be tested by future experiments.

**Figure 4:**
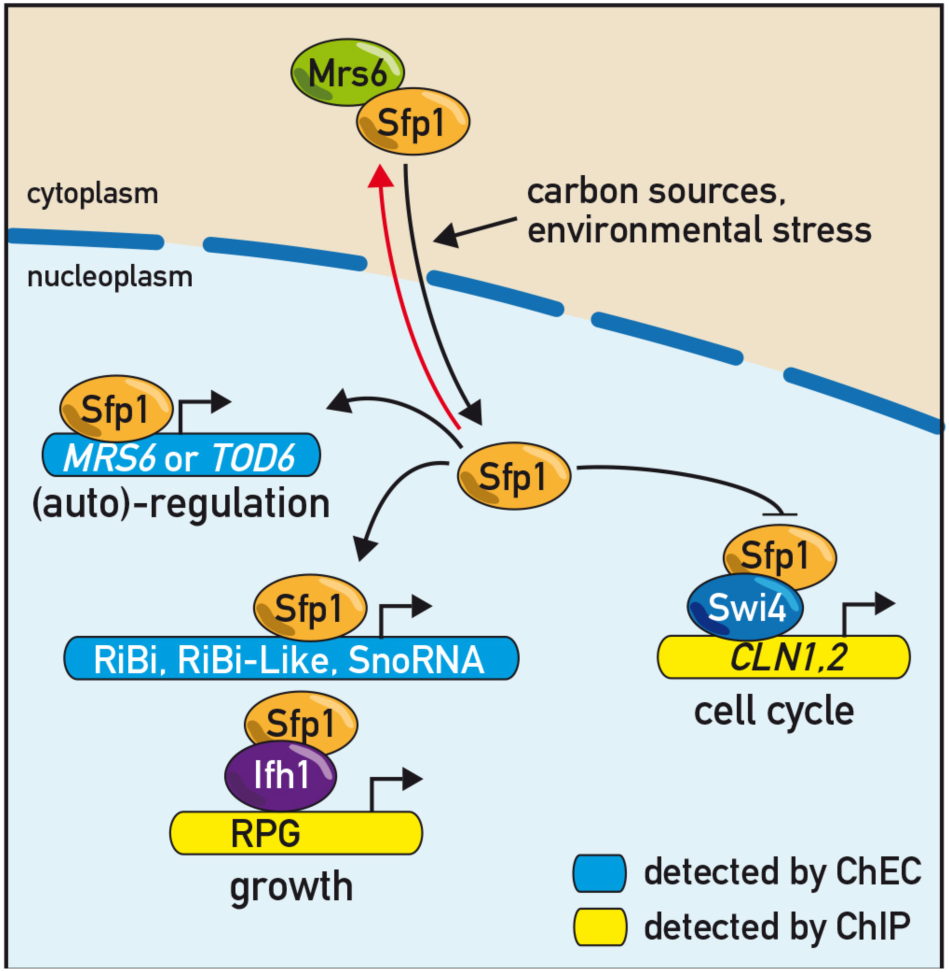
Schematic representation of Sfp1 binding and regulation. According to growth conditions Sfp1 shuttles between the nucleus and the cytoplasm allowing it to adjust TBP and RNAPII recruitment at a vast array of growth-related genes including RiBi, RiBi-like, snoRNA, and RP genes. Sfp1 also binds to and controls expression of its own regulator (*MRS6*) and that of the RiBi gene repressor *TOD6*, possibly to facilitate the rapid shut-down of growth-related transcription upon stress. Finally, Sfp1 is recruited at the promoters of START-specific genes, such as the two G1/S cyclin genes *CLN1* and *CLN2*, in a Swi4-dependent manner. The ability of Sfp1 to bind to and regulate a wide variety of promoter, places Sfp1 an ideal position to co-ordinately regulate cell growth and the commitment to cell division (START) at a transcriptional level.

Although Sfp1 has long been implicated in cell size determination, it has been unclear whether its role is exclusively related to activation of ribosome biogenesis programs or if it also serves as a more direct inhibitor of START (Aldea et al., 2017, Ferrezuelo et al., 2012, Jorgensen & Tyers, 2004). Our identification of *CLN1* and *CLN2* as targets of negative regulation by Sfp1 is supports the latter hypothesis and warrants further study. Although still speculative, we note that inhibition of *CLN1/2* expression by Sfp1 would be expected to delay START and thus prolong growth before division occurs, consistent with the observation that *sfp1Δ* cells are unusually small compared to WT cells. We also note that the glucose-dependent binding of Sfp1 at *CLN1/2* promoters may explain their repression by the cyclic AMP signaling pathway (Baroni, Monti et al., 1994, Tokiwa, Tyers et al., 1994), which is activated by glucose addition to cells growing in poor carbon sources. Nevertheless, the association of Sfp1 with a large number of other genes in the G1/S regulon raises the possibility that Sfp1 regulation of START may extend well beyond its role in *CLN1/2* expression.

The application of ChEC-seq and related MNase-based methods (“Chromatin Immuno-Cleavage [ChIC] (Schmid et al., 2004) or “Cleavage Under Targets and Release Using Nuclease” [CUT&RUN] (Skene & Henikoff, 2015)) is still in its infancy. Nevertheless, we are unaware of other cases, as described here for Sfp1, where ChIP and in vivo MNase-cleavage methods yield such contrasting results. Significantly, both ChIP and ChEC for Sfp1 are concordant with functional data (transcriptional changes upon depletion or over-expression) even though each reports on only a subset of the Sfp1 regulatory landscape. Our findings thus highlight limitations of both techniques for measuring chromatin association of specific proteins, that may be under-appreciated. For example, the failure of ChIP to detect Sfp1 binding at RiBi and RiBi-like genes would appear surprising considering our evidence that these interactions result from direct DNA binding. One possible explanation for this discrepancy is that the proposed Sfp1 binding motif, the RRPE element, is extremely A/T-rich and may thus be unable to form direct cross-links with Sfp1 at a detectable frequency (Rossi, Lai et al., 2018). Alternatively, or in addition, the C-terminal epitope tags so far used to detect Sfp1 by ChIP may be masked at sites where Sfp1 binds directly to DNA, but not at those sites where its binding is dependent upon a second TF. In the case of ChEC, we imagine that Sfp1 detection at RP and G1/S regulon genes might be limited by a short binding half-life and/or access of the tethered MNase to accessible promoter DNA. We suggest that the pleiotropic chromatin-binding behavior of Sfp1 described here is not unique and propose that the complementary application of ChEC-seq and related techniques maybe be essential for identifying the full spectrum of TF targets, not just in yeast, but also in more complex metazoan organisms.

## Methods

### Yeast strains

A complete list of all strains used in this study is provided in the Supplementary Tables 6. Strains were generated by genomic integration of tagging or disruption cassettes (Longtine, McKenzie et al., 1998, Rigaut, Shevchenko et al., 1999).

### Yeast growth conditions

Experiments were performed with log phase cells harvested between OD_600_ 0.4 and 0.6. Yeast strains used in this study are listed in Table 6. Overnight cultures were diluted to OD_600_ = 0.1, grown at 30°C to exponential phase (OD_600_ = 0.4), and then treated with rapamycin at 1μg/ml (from a 1 mM stock solution in 90% ethanol, 10% Tween-20) for anchor-away experiments. Genome-wide localization of Sfp1-TAP was analyzed under standard growth conditions in YP Galactose 2%, Raffinose 2% or Glucose 2%, and the untagged wild type (WT) strain (YDS2) was used as a control. The strain expressing *pGAL1-SFP1-TAP* was grown in raffinose-containing medium for two generations and subsequently treated for 1 hr with 2% galactose to induce *SFP1* expression. For glucose pulse experiments, WT strains were grown in YP glycerol (3%), glucose (0.05%) and shifted to yeast extract, peptone, adenine, and dextrose medium (YPAD; 2% glucose).

### Yeast growth assays

Yeast strains were grown in the appropriate medium to a concentration of 1 × 10^7^ cells/ml. Serial 10-fold dilutions were spotted either on YPAD plates or on plates containing selective medium, at the indicated temperature. Plates were photographed after 2 days of incubation unless otherwise noted.

### Live cell microscopy

All cultures for microscopy experiments were grown to early exponential phase in riboflavin-free medium. Rapamycin was directly added to the cultures at a final concentration of 1 μg/ml. Images were acquired using a wide-field fluorescence microscope (Zeiss Axio Imager Z1m) equipped with a CCD camera.

### ChIP-seq

Cultures of 200 ml were collected at OD_600_ 0.5-0.8 for each condition. The cells were crosslinked with 1% formaldehyde for 10 min at room temperature and quenched by adding 125 mM glycine for 5 min at RT. Cells were washed with ice-cold HBS and resuspended in 3.6 ml of ChIP lysis buffer (50 mM HEPES-Na pH 7.5, 140 mM NaCl, 1mM EDTA, 1% NP-40, 0.1% sodium deoxycholate) supplemented with 1mM PMSF and 1x protease inhibitor cocktail (Roche). Samples were aliquoted in 6 Eppendorf tubes and frozen. After thawing, the cells were broken using Zirconia/Silica beads (BioSpec). Lysates were spun at 13’000 rpm for 30 min at 4°C. The pellet was resuspended in 300 µl ChIP lysis buffer + 1mM PMSF and sonicated for 15 min (30” on -60” off) in the Bioruptor (Diagenode). Sonicated lysates were then spun at 7’000 rpm for 15 min at 4°C. Sfp1-TAP, RNAPII, and TBP-Myc binding were analyzed using TAP-specific, Rpb1, and anti-Myc antibody, respectively (Thermo Fisher CAB1001 or Abcam 5131, Myc epitope 9E10). The antibody (1 μg per 300 μl of lysate) was added to the supernatant and incubated for 1h at 4°C. The magnetic beads were washed three times with PBS plus 0.5% BSA, added to the lysates (30 μl of beads per 300 μl of lysate) and incubated for 2 hr at 4°C. The beads were then washed twice with 50 mM HEPES-Na pH 7.5, 140 mM NaCl, 1mM EDTA, 0.03% SDS, once with AT2 buffer (50 mM HEPES-Na pH 7.5, 1 M NaCl, 1mM EDTA), once with AT3 buffer (20 mM Tris-Cl pH 7.5, 250 mM LiCl, 1mM EDTA, 0.5% NP-40, 0.5% sodium deoxycholate) and twice with TE. Chromatin was eluted from the beads by resuspending in TE + 1% SDS and incubation at 65°C for 10 min. The eluate was transferred to an Eppendorf tube and incubated overnight at 65 °C to reverse the crosslinks. DNA was purified using the High Pure PCR Cleanup Micro Kit (Roche) and libraries were prepared for sequencing using the TruSeq ChIP Sample Preparation Kit (Illumina) according to the manufacturer’s instructions. The libraries were sequenced on a HiSeq 2500 machine and the reads were mapped to the sacCer3 genome assembly using HTSstation (David, Delafontaine et al., 2014).

### Sfp1 binding

ChIP-seq peaks of Sfp1 binding were defined by shifting the plus and minus strand ChIP-seq profiles towards each other by 150 bp and extending each read by 40 bp. To quantify ChIP-seq signals for each promoter, a ratio between the total number of reads from each sample in a 400 bp region upstream the transcription start site (TSS; (Jiang & Pugh, 2009)) of each ORF and the total number of reads from the same region obtained with mock IP of the control untagged strain. The same logic was applied to quantify signals within ORFs.

### Swi4 and Ifh1 binding

ChIP data from Harbison et al. (2004) and Knight et al. (2014) were used to map Swi4 and Ifh1 binding, respectively. The ChIP-seq peaks (Knight et al., 2014) were defined by shifting the plus and minus strand ChIP-seq profiles towards each other by 150 bp and extending each read by 40 bp. To quantify ChIP-seq signals for each promoter, the total number of reads from each sample in a 400 bp region upstream the TSS (transcription start site; (Jiang & Pugh, 2009)) of each ORF was determined.

### TBP binding

ChIP-seq signals for TBP were quantified at (TBP binding site) positions taken from (Rhee & Pugh, 2012).

### Rpb1 (RNAPII) binding

To quantify Rpb1 ChIP-seq signals for each gene, a ratio was calculated of the total number of reads in each ORF before treatment to the total number of reads in each ORF after the indicated times of rapamycin or vehicle treatment, or after 1h in galactose for the strain carrying *pGAL1-SFP1-TAP*. In the Sfp1-FRB anchor-away experiment measuring Rbp1 ChIP, *S. pombe* chromatin was used as a “spike-in” control for normalization, as described previously (Bruzzone et al., 2018).

### ChEC-seq

ChEC-seq experiments were performed essentially as described (Zentner et al., 2015) with the following modifications. Cells in which MNase was fused at the C-terminus of the endogenous *SFP1* gene were used to determine Sfp1 binding. Cells in which MNase was placed under the control of *REB1* promoter were used as a control. One sample corresponds to 12 ml of culture at OD_600_ = 0.7. Cells were washed twice with buffer A (15 mM Tris 7.5, 80 mM KCl, 0.1 mM EGTA, 0.2 mM spermine, 0.5 mM spermidine, 1xRoche EDTA-free mini protease inhibitors, 1 mM PMSF) and resuspended in 200 μl of buffer A with 0.1% digitonin. The cells were incubated for 5 min at 30°C at which point MNase was induced by addition of 5 mM CaCl_2_ and stopped at the desired timepoint by adding EGTA to a final concentration of 50 mM. DNA was purified using MasterPure Yeast DNA purification Kit (Epicentre) according to the manufacturer’s instruction. Large DNA fragments were removed by a 5-min incubation with 2.5x volume of AMPure beads (Agencourt) after which the supernatant was kept, and MNase-digested DNA was precipitated using isopropanol. Libraries were prepared using NEBNext kit (New England Biolabs) according to the manufacturer’s instructions. Before the PCR amplification of the libraries small DNA fragments were selected by a 5-minute incubation with 0.9x volume of the AMPure beads after which the supernatant was kept and incubated with the same volume of beads as before for another 5 min. After washing the beads with 80% ethanol the DNA was eluted with 0.1x TE and PCR was performed. Adaptor dimers were removed by a 5-min incubation with 0.8x volume of the AMPure beads after which the supernatant was kept and incubated with 0.3x volume of the beads. The beads were then washed twice with 80% ethanol and DNA was eluted using 0.1x TE. The quality of the libraries was verified by running an aliquot on a 2% agarose gel. Libraries were sequenced using a HiSeq 2500 machine in single-end mode. Reads were extended by the read length. To analyze the Sfp1-MNase binding pattern, read ends were considered to be MNase cuts and were mapped to the genome (sacCer3 assembly) using HTSstation (David et al., 2014). For peak analysis MACS software was used through HTSstation, using free-MNase signal as background. Motifs were detected using MEME (Bailey, Boden et al., 2009) with sequences from each identified ChEC signal peak as input.

## Data and software availability

All sequencing and microarray data generated in this study were submitted to the GEO database as SuperSeries GSE118561.

## Acknowledgements

We thank other members of the Shore laboratory for helpful discussions; Florian Steiner and Robbie Loewith for comments on the manuscript; Uli Laemmli for ChEC reagents and advice on the ChEC method; Mylène Docquier and the Institute of Genetics and Genomics of Geneva (iGE3; http://www.ige3.unige.ch/genomics-platform.php) for high-throughput DNA sequencing; Nicolas Roggli for expert artwork; and Thomas Schalch for the use of his local Galaxy server. B.A. acknowledges support from an EMBO Long-Term Fellowship. D.S. acknowledges support from the Swiss National Fund (grant number 31003A_170153) and the Republic and Canton of Geneva.

## Author contributions

B.A. and S.T. designed the study, together with D.S., and carried out most of the experiments. Y.G. constructed and characterized Sfp1 anchor-away strains, B.A. and S.M. performed all ChEC-seq experiments, and S.K. analysed RNAPII binding on coding regions and the “spike-in” control for RNAPII ChIP-seq following Sfp1 depletion. B.A. and D.S. wrote the manuscript.

